# Correcting bias from stochastic insert size in read pair data — applications to structural variation detection and genome assembly

**DOI:** 10.1101/023929

**Authors:** Kristoffer Sahlin, Mattias Frånberg, Lars Arvestad

## Abstract

Insert size distributions from paired read protocols are used for inference in bioinformatic applications such as genome assembly and structural variation detection. However, many of the models that are being used are subject to bias. This bias arises when we assume that all insert sizes within a distribution are equally likely to be observed, when in fact, size matters. These systematic errors exist in popular software even when the assumptions made about data are true. We have previously shown that bias occurs for scaffolders in genome assembly. Here, we generalize the theory and demonstrate that it is applicable in other contexts. We provide examples of bias in state-of the-art software and improve them using our model. One key application of our theory is structural variation detection using read pairs. We show that an incorrect null-hypothesis is commonly used in popular tools and can be corrected using our theory. Furthermore, we approximate the smallest size of indels that are possible to discover given an insert size distribution. Two other applications are inference of insert size distribution on *de novo* genome assemblies and error correction of genome assemblies using mated reads. Our theory is implemented in a tool called GetDistr (https://github.com/ksahlin/GetDistr).

## 2 Introduction

Insert size distribution When sequencing paired reads (*e.g.* paired end sequencing, mate pair sequencing, fosmid or BAC ends [1, 24]), one targets a specific *insert size* (the genomic distance between the *reads* in a read pair). The insert size is dependent on the aims of the project. For example, genome assembly can benefit from using libraries with a range of insert sizes [16, 26]. Another example is detection of structural variants. Longer insert sizes improves span coverage which yields more observations over a variant and they also span longer insertions. On the other hand, shorter insert size libraries tend to have less variation around its mean which facilitates more accurate predictions.

Although targeting an insert size when sequencing paired reads, the insert size of fragments varies. This variation gives an uncertainty that can expressed as a probability distribution. We will have a mean insert size *μ* and a standard deviation *σ* around its mean. This probability distribution is discrete since it is given in number of base pairs. In some cases, one can approximate the insert size distribution with a normal distribution (a common assumption in various tools, [18, 15, 4, 8, 14]), going from discrete to continuous support. This approximation is in many cases not accurate as insert size distributions often have more observations far out from the mean (thicker tails) while some distributions deviates strongly from a normal distribution on most of it’s support [16].

### Truncated and skewed insert size distribution

Inferring insert size distribution is easy if we can observe all insert sizes from a read pair library. In that case, all insert sizes contribute with equal “weight” when calculating *e.g. μ* and *σ*. In many applications we do not observe the insert size of all read pairs, for example, when inferring the insert size distribution of paired reads spanning over a region of *δ* base pairs. The insert size distribution of the spanning read pairs will not follow the same distribution as the insert size distribution of the whole library. This is seen from the fact that read pairs with insert size smaller than *δ* base pairs will not span over the region. But this is not the only reason to why the observed distribution is different; longer sequences have more positions from where it can span over the region. Thus, in terms of probability, it is more likely for longer read pairs to span over *δ* base pairs. The probability for given insert size to span a region is not only determined by the insert size distribution. It also depends on the length of the insert size compared to the region it is supposed to span. We can think of the latter criterion as the “weight” to observe a given read pair from the library distribution. The weight is given by the number of possible placements a read pair of given insert size can have [19].

### Our contributions

We state a generative model (DRISM) and derive probabilities for insert size parameters in different scenarios. Using these probabilities, we derive more accurate mate pair library insert size estimations on fragmented (*de novo*) assemblies. We correct a commonly assumed null-hypothesis for detecting structural variants with insert size based methods. We also show how the corrected null-hypothesis evens out the detection of insertions to deletions for the insert size based variant caller CLEVER [15]. Furthermore, we improve size prediction of insertions for CLEVER and BreakDancer [3]. We also study what size of indels that are possible to discover given DRISM. The theory presented in this work also generalizes to other applications using insert size based inference such as genome assembly evaluation with paired reads [6, 10, 23].

## 3 Model

We now state the Distribution of Read Insert Sizes Model (DRISM), based on the Lander-Waterman model [11], which we use for deriving probabilities in section 4. We will refer to *target sequence* as a DNA fragment on which read pairs are aligned, *e.g.*, a genome, contig or exon. DRISM assumes that read pairs are sampled randomly with a uniform distribution over the genome, giving a Poisson distributed coverage (*i.e.*, the Lander-Waterman model). Let *G* denote the length of the genome (estimated or known). Data for the model comes from alignment of reads where we observe: distance *o* between reads in a read pair, read length *r*, number of allowed softclipped bases [13] in an alignment *s*, the length of target sequence(s) *a* (and b), see Figure 1. Furthermore, DRISM assumes that reads pairs comes from some library distribution *f*(*x*) where *f*(*x*) may or may not be known, dependent on application (see section 3). Finally, a parameter *δ* models the number of base pairs that is unobserved between two reads in a pair. For example, *δ* models the gap between two contigs in *de novo* assembly or the size of an insertion/deletion in structural variant detection. *δ ∈ Z* where negative values represent overlapping sequence while positive values represent a missing sequence. We will discuss the model in two different settings.

**Figure 1:**
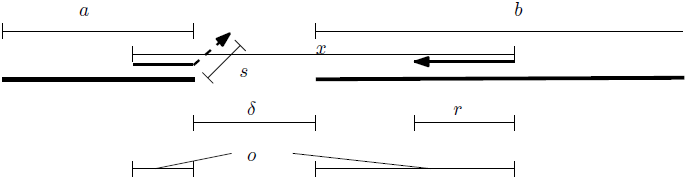
Constant and variable notations in DRISM.

- *Complete target sequence:* Inference of *μ* and *σ* (*e.g.* for estimation of mate pair distributions in *de novo* assembly, the target sequences are the contigs with known length, hence *δ* = 0).
- *Incomplete target sequence:* Inference of *μ* and *σ* of the truncated and skewed insert size distribution (*e.g.* for detection or size estimation of structural variants, or size estimation of gaps between contigs in *de novo* assembly).

These scenarios are two examples of when DRISM is useful. Having an explicitly formulated generative model will aid deriving probabilities where reads are sequenced from a genome.

## 4 Deriving probabilities from DRISM

### 4.1 Distribution over complete target sequence

We assume that the target sequence is known and therefore *δ* = 0. We will derive the probability that we observe a read pair with insert size of length *x* placed on a target sequence of length *a* coming from a genome of length *G*. Let *f*(*x*) be the insert size distribution and *w*(*x*) be the number of possible placements of a read pair with insert size *x* onto *a*. We then write

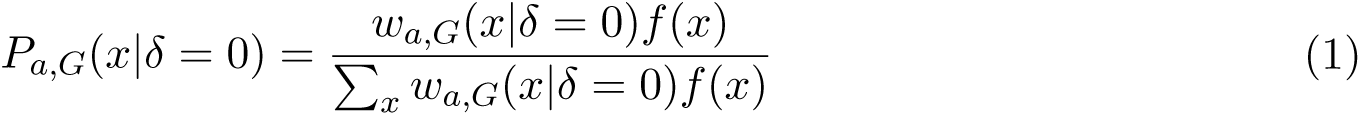

where *P*_*a,G*_(*x|δ* = 0) is the probability to observe an insert size *x* on a target sequence of length *a* from a genome of length *G*. The denominator is a normalization constant to make *P* a probability. Subscript on *a, G* is used instead of conditions in functions (*e.g.*, *P*_*a,G*_(*x*) instead of *P* (*x|a, G*)) due to readability. Note that *f* is independent of *a* and *G*. We have that *w*_*a,G*_ (*x*) can be written as

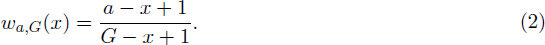

For example, 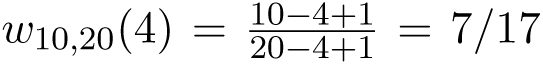, which states that a read pair of insert size 4 will have 7 possible placements on a target sequence of length 10 (the nominator). It also has 17 possible placements on a genome of length 20 (denominator). This together gives a probability of 7*/*17 that it places within a target sequence of length *a*.

#### Softclipping

Reads can be partially aligned to a target sequence with the remaining part “hanging off the end”. The number of bases that falls outside the target is referred to as *softclipped* bases [13]. To allow for softclipped bases, we introduce an additional parameter *s* denoting how many base pairs are allowed to be softclipped. We get *wa*(*x*) as

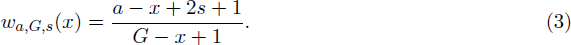

For example, 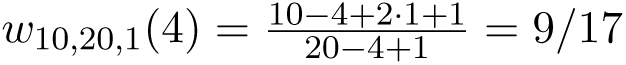. The probability has now increased compared to (2) since the read pair is allowed two more placements (we can slide it outside the target sequence one extra position to the left, and one to the right). Notice that *s* is not introduced in the denominator since read pairs are assumed to come from the genome.

#### Simplifications

From now on, we will let the denominator of (3) be approximated by the constant *G*. This is motivated by the small variation effect of *x* has on the number *G − x* +1, (*i.e.*, in practice *G − x* + 1 *≈ G*). This will make calculations of likelihood functions easier. We will not consider the special case where a target sequence is the very end of a genome (in case of a linear genome), which will implicate that we cannot have softclipped bases on one end of the target sequence. In that particular case, it would slightly change the nominator (3). The effect of these two modifications on the probabilities is however vanishingly small. Thus, we write

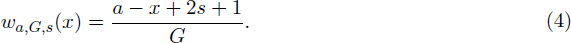

#### Several target sequences

In case we have *u* target sequences, *e.g.* contigs in a genome assembly, we need to generalize the weight function (3) to give the probability of observing a read pair with insert size *x* on any of the target sequences. This is done by summing over all possible positions on the target sequences of lenghts *ai*, *i* = 1, *…, u* as

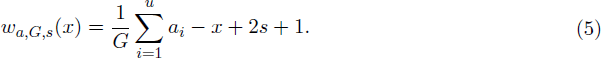

### 4.2 Distribution over incomplete target sequence

In this section we will discuss the case when *δ* is unknown. There are cases where we can not observe true insert size of observed reads, they might be located on different target sequences, *e.g.,* in contig scaffolding or they might span a gap of unknown length on the target sequence (*e.g.* over a structural variation. We want to find the probability to observe a read pair with insert size *x* over a given gap *δ* and given target sequence(s) with length(s) *a* (and *b*). We will make use of the following relation *δ* = *x − o* (see Figure1). To give a probability for this scenario, we proceed to describe the adjustments to the probability in model (1).

#### Modifying *f*(*x*)

The stochastic variable *X* is now unknown since we can not observe *δ*. The distribution of *f* will not change since it is the full insert size distribution. However, we change the variable to write *f*(*o* + *δ*).

#### Modifying *w*(*x, δ*)

With the simplifications of the denominator in section 4.1, *w* can be regarded as a weight function that indicates how many placements a read pair can have over *δ* since the denominator is fixed for any *x*. In the case of one target sequence, there are two boundaries on the number of placements. The read pair needs to be larger than *δ* and have sufficiently many base pairs on each side to be mapped (lower boundary). It also needs to be shorter than *a* (upper boundary). We have

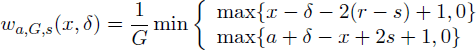

The 0’s in the max function are there to keep the function weights to 0 in case we have no possible placing of a paired read. However, we can simplify this function to be expressed in *o*, which is known, instead of *x* and *δ*. We have

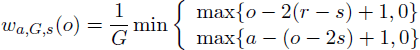

If there are two different target sequences,*e.g.* two contigs, the number of placements can be limited in three different ways [19]. We have the two boundaries above, plus the case where one of the target sequences are so small that it restricts the positions of all paired reads that spans the two target sequences.

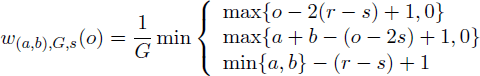

#### Probability for *δ*

For clarity, we will from now on omit the subscripts over *a,* (*b*), *G, s*. We assume that we have no information of *δ* beforehand, thus we let the distribution over *δ* be uniform. We can express the probability for *δ* given observations as

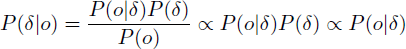

The first proportionality comes from the fact that *o* is known which makes *P* (*o*) a constant and the second because we assume a uniform prior for *P* (*δ*). We now have

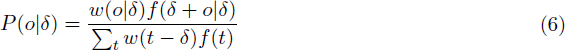

Where the denominator is the sum of all possible insert sizes that can be observed given *δ* and *f*. A practical interval for *t* would be *t ∈* {*δ* + 2(*r − s*), *μ* + 6*σ*}. We can now find the most likely *δ* using maximum likelihood estimation (MLE) over (6). Note that we implicitly get *P* (*x|δ*) since

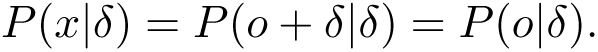

Also note that we have *G* as a constant denominator in *w* and *w* occurs both in the nominator and denominator of (6), therefore it can be canceled out when computing the MLE.

#### Naive estimation of *δ*

Many tools that estimates unknown sequence length (*e.g.* in assembly and structural variation detection) by observed insert sizes use the following formula

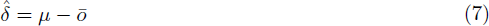

where *ō* is the mean observation [3, 15, 7, 21, 4, 2, 5]. At first glance this formula seems reasonable since we take the expected insert size and subtract the mean of the observations. However, this formula has strong limitations. One of the most obvious is that 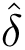 has an upper bound of *μ* − 2(*r* − *s*) in this case (since *o ≥* 2(*r − s*)). This equation implies that we can never span over a sequence longer than *μ −* 2(*r − s*). This is of course not correct.

## 5 Applications and Results

The following section shows how the probabilities derived in (1) and (6) can be used to improve inference with read pairs. We compare our results against some popular state-of-the-art algorithms in their respective fields. *We want to stress that it is our aim to illustrate examples of biases in popular tools. We do not infer general performances of the tools we use in the evaluation.* The software tested here are some of the most popular and best suited for their respective tasks. Making inference with an exact insert size distribution may or may not be a of high importance in their respective applications. Nevertheless, being aware of biases that occur improves analysis of results. Also, it is important to be aware of these biases for future method development. Here, we show how to apply our theory on two applications (other than scaffolding as was shown in [19]).

We have performed our experiments under ideal circumstances, and show that there exist biases due to model errors. That is, we simulate data that follows the assumptions of the tools we test and are still able to show that systematic errors occur. It is not our intention to give a model that handles alignment errors or library preparation biases. However, the authors believe that it is possible to extend the theory for this and it is subject for future work.

### 5.1 Library insert size estimation from alignment

In *de novo* assembly projects, data sets are often obtained at different time points since it is not always clear at first what data is needed to assemble a genome and also, technology might improve during the period of a project. Draft assemblies are created while new read libraries are obtained. In order to make good use of additional libraries, we want to obtain correct read library metrics. Inference of these metrics without a reference genome is difficult and a common procedure is to map the new library to a draft assembly (also, this mapping can reveal erroneous regions in the assembly). We will here discuss issues with inference of read library metrics in *de novo* assembly.

We compared insert size estimation on a mate pair library from Rhodobacter sphaeroides [17] using picard [9], BWA [12] and GetDistr. BWA and picard use a pre filtering step when estimating *μ* and *σ*. BWA uses reads that have a PHRED mapping quality *≥* 20 and insert size within [*Q*1 *−* 2(*Q*3 *− Q*1), *Q*3 + 2(*Q*3 *− Q*1)] where *Q*1, *Q*3 are the first respectively third estimated quartile. Picard selects reads between 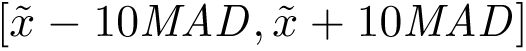, where 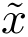 is the median and *MAD* is the median absolute deviation. For this data set, the lower and upper boundary of insert sizes used for each tool were approximately [0, 6500] for BWA and [0, 11000] for picard. GetDistr was used with both of the heuristic filters from BWA and picard. We first estimated *μ* and *σ* on the reference genome of Rhodobacter sphaeroides using all tools and then compared these to estimates obtained by mapping the reads to contigs of the Allpaths-LG assembly from GAGE [22]. Figure 2a and 2b illustrates the results where x-axis starts with estimates on the reference sequence followed by estimates from mapping the read library to contigs with size *≤ N x* of the assembly. That is, *x* = 0 corresponds to the full Allpaths-LG assembly, *x* = 10 corresponds to the remaining assembly after removing contigs with size *≤ N* 10, etc. GetDist has a subscript denoting which heuristic that was used to select reads. As seen in figure 2a and 2b, the filtering does not affect GetDistr estimates much. Since picard and GetDistr agrees on roughly 2600 as mean insert size and 1325 as standard deviation, this is believed to be close to the truth. Even though the assembly is of very high quality and contiguity compared to the insert size (N50 of 42.5kbp), we see that the estimates are affected (*e.g.* shifted downwards for the mean). The effect becomes clearer as estimates are obtained from smaller contigs (as *x* increases). GetDistr is unaffected by this until *x* = 80, when most contigs are smaller than the true mean insert size. The same method can be used to adjust *e.g.* the median and *MAD* of the library as well.

**Figure 2:**
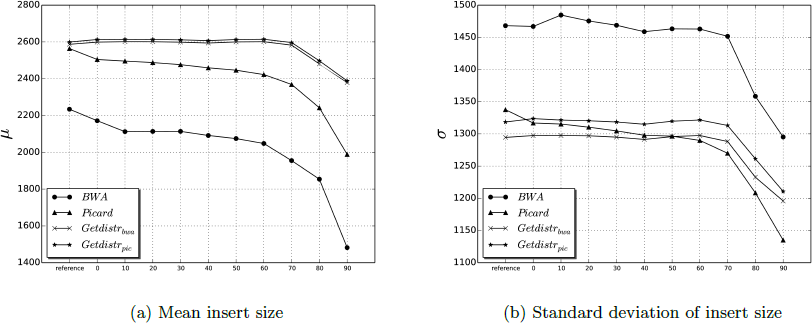
Estimation of library mean and standard deviation for a mate pair library of R. sphaeroides. Input to GetDistr and picard is from mapping reads with BWA on the reference sequence (leftmost point on x-axis) and different contig sets from the Allpaths-LG assembly (numbers on x-axis). The number on the x-axis denotes that contigs with size less than *N x* are used for inferring mean and standard deviation. Library estimations decreases in quality as shorter contigs are used. Our probabilities accounts for contig lengths and gives more accurate and consistent estimations across different lengths of contigs.

### 5.2 Structural variation

Under the null-hypothesis, no variant is assumed, *i.e. δ* = 0, thus *x* = *o*. Let 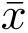 denote the average insert size observed over a position *p* on the reference sequence and, as before, *μ* be the mean of the library insert size distribution *f*. Assuming the DRISM model with *f* normally distributed, a commonly used null-hypothesis is then *H*_0_ : 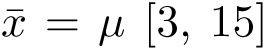, and statistical testing of variants are performed using *H*_0_. That is, a test showing if the 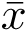 deviates significantly from *μ* over *p*. We claim that this null-hypothesis is wrong because *μ* is not the expected mean over *p*. Using the theory in section 4.2 we have

**Result1.** *Given DRISM, f ∼ N* (*μ, σ*), *we have 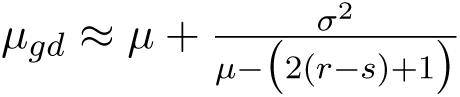*

**Result2.** *Given DRISM, f ∼ N* (*μ, σ*), *we have 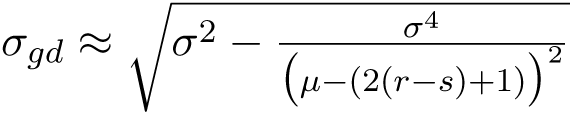*

See appendix A.3 for derivation. From a theoretical point of view, we can also derive an approximate formula for the smallest size of indels we can expect to find with insert size based methods. We let the smallest indel size be denoted by *m* and coverage be denoted by *c*. Given parameters (*μ, σ, r, s, c*) and *f ∼ N*, we want to find the smallest size of an indel *m* that we can detect given the DRISM model and a controlled of false positive rate *α*. As real data introduce noise, *m* can be seen as a lower boundary. We can state this problem as finding the smallest value *m* such that the probability below holds.

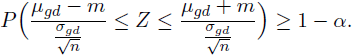

Note that the smallest *m* will give an equality sign in the equation above. From this equation, we get

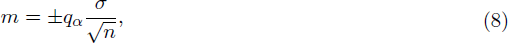

where *q_α_* denotes the quantile value at *α*. Since this formula depends on the number of observations, it is base pair specific. To get an approximate estimate of *m* across the genome, we need to find the expected number of observations *n* over a base pair. We get this estimate similarly to [25] with the modification of adding parameters *r* and *s* to their formula. Table 1 shows some examples of *m* given *α* = 0.05 and the Bonferroni correction to control the family-wise error rate. Considering that significance tests are correlated in this context, Bonferroni correction is conservative for these tests. Due to the approximations and assumptions used in this derivation we not place too much weight on analyzing the exact numbers in the tables. However, it is important to note the effect of these parameters on *m*. Not only is *σ* and *c* important as reported [15, 3], *μ* contributes a substantial amount to the quality as it increases *n* for each test (higher span coverage).

**Table 1:** Approximate values of *m* given given *G* = 3 · 10^−9^, *α* = 0.05 *r* = 100 and *s* = 0. With a two sided test and Bonferroni correction, a significant p-value lies outside 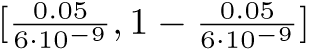.

| $\mu$ | | 300 | | | 400 | | | 500 | | |
| --- | --- | --- | --- | --- | --- | --- | --- | --- | --- | --- |
| $\sigma$ | | 25 | 50 | 75 | 25 | 50 | 75 | 25 | 50 | 75 |
| $c$ | 20 | 53.2 | 106.2 | 157.2 | 37.6 | 75.3 | 112.9 | 30.7 | 61.5 | 92.2 |
|  | 50 | 33.7 | 67.2 | 99.4 | 23.8 | 47.6 | 71.4 | 19.4 | 38.9 | 58.3 |
|  | 100 | 23.8 | 47.5 | 70.3 | 16.8 | 33.7 | 50.4 | 13.7 | 27.5 | 41.2 |

#### Size estimation of insertions

We compared estimation of insertions of two insert size based structural variation tools, BreakDancer [3] and CLEVER [15]. From a simulated reference genome, we generated insertions in a donor genome and a paired read library with *f ∼ N* (500, *σ*), *σ ∈* [25, 50, 75, 100], *r* = 100 and *c* = 100 (*f ∼ N* is assumed in CLEVER and BreakDancer). 100 insertions with size *i* where generated where *i* = 25, 50, 100, 150, 200, 250, 300, 350. We aligned the reads back to the reference genome with BWA [12].

Panel 1 and 2 in figure 3 show GetDistr predictions vs naive predictions (using equation 7) of insertion size. These predictions were obtained from all reads spanning over the insertions. Panel 3-6 show predictions from BreakDancer and CLEVER, together with predictions by GetDistr from the same set of reads as in the tools. Similarly to [15], a prediction is classified as a true positive, hence included in the size analysis, if the break point prediction is within the internal segment size base pairs away from the true breakpoint (*i.e.* at most *μ −* 2*r* base pairs away). BreakDancer gives a confidence interval for its predictions. We therefore used the median point as inferred break point position. We see that without changing the methodology for detecting variants in the tools, we are able to improve predictions using our theory. Because of different methodologies in BreakDancer and CLEVER, the biases are different. BreakDancer uses only reads with 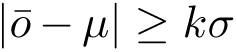 (with *k* = 3 default). CLEVER uses more sophisticated statistical testing, taking all read pairs into account in a first step, but uses only subsets of them to predict the variant. It should be noted that even though GetDistrs probabilities are not tailor made for BreakDancers and CLEVERs heuristics (*i.e.*, using subsets of reads with special properties), we still improve the predictions considerably. When using insert size for variant detection, our recommendation is to not restrict the analysis only to anomalous read pairs. Also note that the number of insertions detected are highly dependent on *i* and *σ*. Some points in Figure 3 contains few predictions (see Table 2).

**Figure 3:**
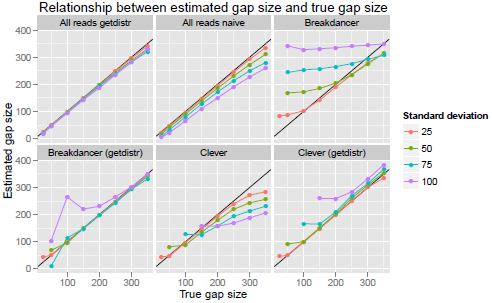
Length estimates of insertions on a simulated genome.

**Table 2:**
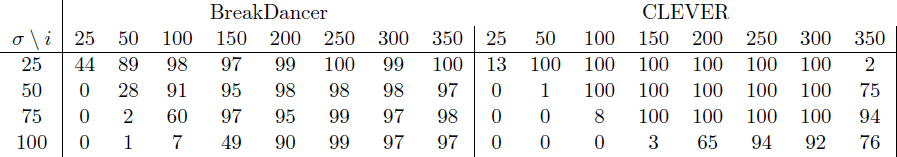
Number of insertions detected by CLEVER and BreakDancer for each point in figure 3.

| $\sigma \setminus i$ | BreakDancer | | | | | | | | CLEVER | | | | | | | |
| --- | --- | --- | --- | --- | --- | --- | --- | --- | --- | --- | --- | --- | --- | --- | --- | --- |
|  | 25 | 50 | 100 | 150 | 200 | 250 | 300 | 350 | 25 | 50 | 100 | 150 | 200 | 250 | 300 | 350 |
| 25 | 44 | 89 | 98 | 97 | 99 | 100 | 99 | 100 | 13 | 100 | 100 | 100 | 100 | 100 | 100 | 2 |
| 50 | 0 | 28 | 91 | 95 | 98 | 98 | 98 | 97 | 0 | 1 | 100 | 100 | 100 | 100 | 100 | 75 |
| 75 | 0 | 2 | 60 | 97 | 95 | 99 | 97 | 98 | 0 | 0 | 8 | 100 | 100 | 100 | 100 | 94 |
| 100 | 0 | 1 | 7 | 49 | 90 | 99 | 97 | 97 | 0 | 0 | 0 | 3 | 65 | 94 | 92 | 76 |

#### Detection of indels

In this section we illustrate how the corrected hypothesis 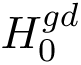 can even out the ratio between detected insertions and deletions. We simulated 100 insertions and deletions respectively with sizes 10, 20, 30, 40, 50, 75, 100. We also simulated three different paired end libraries with *μ* = 300, 400, 500 and *σ ∈* 25, 50, 75. All variations were on a distance of *μ* + 6*σ* from each other and enough reads were generated so that CLEVER estimated *μ* and *σ* within 0.5 base pairs accuracy in all experiments to give perfect conditions. We ran CLEVER as default (*H*_0_) and with 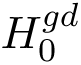 by inserting the corrected values into CLEVER’s code. The results are shown in figure 4. We see that the method used by CLEVER detects more deletions than insertions of the same sizes. Using 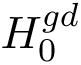, we can reduce this bias to some extent. That is, in general, 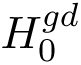 reduces the number of deletions but increases the number of insertions detected. We also reduce the overall false positive rate.

**Figure 4:**
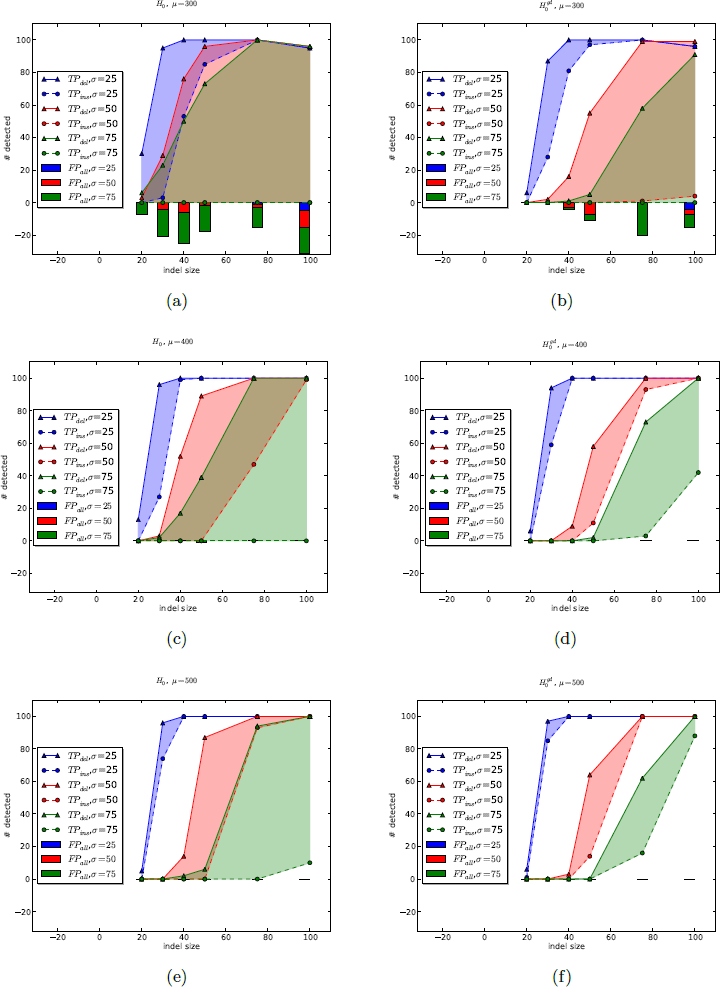
True and false positives for CLEVER when detecting insertions and deletions on a simulated genome with 100x (uniform) coverage from a normally distributed read pair library *N* (*μ, σ*), according to model assumptions by CLEVER. The reference insert size distribution was correctly estimated in all cases. Shaded area displays difference between number of insertions and deletions detected. Bars on negative *y*-axis shows the number of false positives.

## 6 Conclusions

We have derived probabilities from a commonly assumed read pair model which we referred to as DRISM. The probabilities are useful for any method making inference based on read pair libraries. We use the probabilities in different settings to improve predictions of state-of-the-art software. For instance, we derive a more accurate distribution under the null-hypothesis for variant callers. Furthermore, the authors believe that including information such as GC-bias and alignment probabilities in DRISM can be done.

## A Appendix

### A.1 Conditional mean and standard deviation of insert size

Using the DRSIM model with the probabilities we have derived, we can now get theoretical estimates of the expected insert size over an sequence of length *δ*. In figure 5a, we can see how the expected insert size *E*[*X*], varies with *δ* and *σ* if *f ∼ N* (*μ, σ*) and the reference(s) is longer than most of the insert sizes (*e.g. ≥ μ* + 4*σ* if *f ≈ N*). Naive estimation would be *E*[*X*] = 500 independent of *δ, σ*.

**Figure 5:**
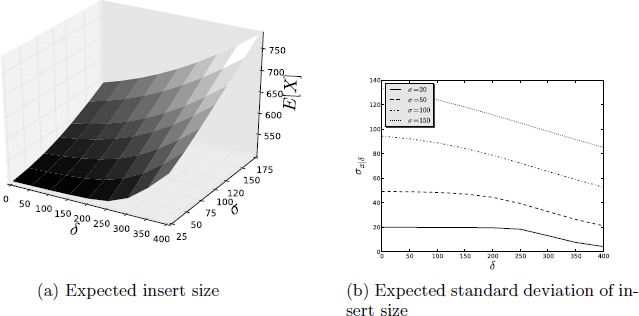
We used *f ∼ N* (500, *σ*) *r* = 100 and *s* = 0 in both plots. **a)** Expected insert size as *σ* and *δ* varies. The mean insert size will deviate more from the full insert size distribution mean *μ* as *δ* and *σ* increases. **b)** Expected standard deviation of insert size over different *δ*. The expected standard deviation of insert sizes *σx|z* will deviate more from the standard deviation of the full insert size distribution *σ* as *δ* and *σ* increases.

The expected value fdfdwqwrwcan be calculated as

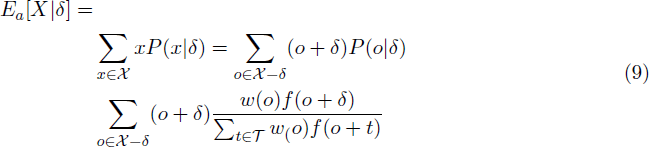

Here, *Τ* is the domain of all possible unknown sequence lengths *δ* given *o*. Figure 5b illustrates how expected standard deviation decreases as *δ* increases. This is expected since a smaller range of insert sizes are able to span a gap of *δ* base pairs as *δ* becomes larger, this is also discussed in [20].

### A.2 Computational complexty

Let 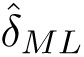 be the maximum likelihood estimate of *δ* and let *m* be the length of the interval where we search for 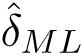 in. Without loss of generality, we can assume that we search for 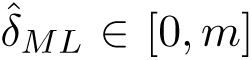 (*m* depends on the full library distribution and can be set heuristically). Let *n* as before be the number of links. For each *δ ∈* [0, *m*] (i.e. *m* times), we need to do *n* calculations in the nominator and *m − δ* calculations in the denominator. *m − δ* will go from *m* to 0 but the highest complexity is reached when we sum up *m* values. This gives the complexity *O*(*m*(*n* + *m*)). From experience, usually *m* dominates *n*, thus we can write *O*(*m*^2^). This is the complexity for brute force calculation of the likelihood function when convexity of the likelihood function cannot be guaranteed. If we can assume convexity, the computational complexity decreases to *O*(*m* log *m*) as binary search or Newton-Raphson algorithm can be employed. Furthermore, if *f*(*x*) follows a normal distribution, [19] showed that the ML value can be obtained with *O*(log *m*) complexity using an analytic expression of the maximum likelihood equation.

### A.3 Insert size estimation − simulated data

In this section we compare our model to BWA’s insert size estimation. This experiment is conducted as follows. Mate pair reads with distribution *N* (*μ, σ*^2^) are uniformly simulated from a repeat free genome. A reference sequence of length *a* is extracted from this genome (*e.g.* a contig). BWA is used as read aligner to map the mate pair reads to the shorter sequence. BWA’s estimate of the parameters *μ* and *σ* is obtained by parsing it’s output. GetDistr’s estimation is obtained from the SAM file produced by BWA. To avoid external effects GetDistr uses only mate pairs with the same characteristics as the mate pairs selected for insert size estimation by BWA. That is, they must have a PHRED mapping quality *≥* 20 and insert size within [*Q*1 *−* 2(*Q*3 *− Q*1), *Q*3 + 2(*Q*3 *− Q*1)] where *Q*1, *Q*3 are the first respectively third estimated quartile. For each combination of *μ*, *σ* and *a*, 100 experiments were run. The results are illustrated in figure 6a and 6b.

**Figure 6:**
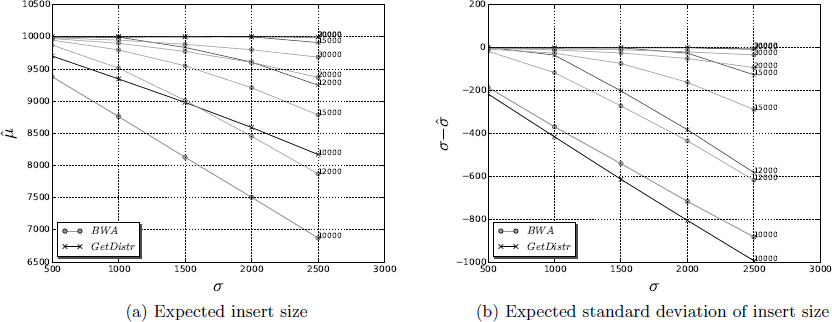
Estimations of library mean and standard deviation. Read library is simulated from *N* (10000, *σ*), *σ ∈* {500, 1000, 1500, 2000, 2500} with *r* = 100. Sequencing depth is 100x coverage. A single line corresponds to different *σ* for one fixed reference length (marker at the end of each line). Reference lengths are 10000, 12000, 15000, 20000, 30000 base pairs. **a)** Estimations of *μ* (y-axis) for different *σ* (x-axis). GetDistr has most of its estimations tightly clustered around the line *μ* = 10000 which is the true mean. **b)** Estimations of difference between true and estimated standard deviation (y-axis) for different library variations (x-axis). GetDistr performs well as long as not a significant part of the library distribution cannot be observed.

Estimations become more biased with shorter reference sequence and higher library standard deviation. GetDistr obtains unbiased estimations of *μ* for all reference length and standard deviation combinations down to 15 000 (figure 6a). It also gives accurate predictions of *μ* even though a significant part of the insert size library cannot be observed (*μ* = 1000, 12000). BWA obtains reasonable estimations only when reference length is 30 000 base pairs. For estimation of *σ*, we are performing better than BWA for reference sequences down to 15 000 base pairs. When *a* = 12000 and 10000, a large part of the distribution falls outside the reference sequence and therefore, an underestimation of *σ* is inevitable. For example, when *a* = 10000, GetDistr, estimates *σ* to almost half of the true standard deviation. This makes sense since only the lower half of the distribution, hence half of the variance, is observed. BWA’s slightly better estimations of *σ* in these cases are a consequence of the underestimated 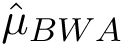.

### A.4 Derivation of Result 1 and 2

From [19] we have 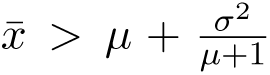, since 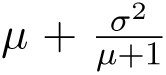 is the expected insert size over any base pair. The greater sign comes from the lack of the constraint that at least *r − s* bases should be aligned on each side of *p*. Such constraint is needed in practice. For example, CLEVER uses *s* = 2 in its implementation which means that at least *r −* 2 base pairs from both reads must be located on respective sides of the variation. BreakDancer has no such criterion, but the criterion is then imposed on the read aligner being able to map at least *r − s* bases on respective sides. This gives the condition *x ≥* 2(*r − s*).

Let *μ_gd_* denote the mean of the distribution of reads spanning *p*. An exact value of *μ_gd_* can be obtained for arbitrary distributions *f* by calculating the expected insert size in equation 9 with *δ* = 0, *a* = *G* and *x ≥* 2(*r − s*). We can however give an accurate approximation of *μ_gd_* by letting *q* = 2(*r − s*) + 1 and substituting the 1’s to *q*’s in [19] (section 2.4, derivation of equation 2). We get Result 1 from this calculation. The derivation is identical, we therefore omit it here and only discuss why it’s an accurate approximation.

The approximation is motivated as follows. The derivation in [19] (section 2.4) is assuming infinite support. Therefore, the above approximation is only accurate if the upper and lower boundaries are far not located near high density regions of *f*(*e.g.*. near the mode if *f ∼ N*). It is easy to motivate that *G* (the upper boundary) satisfies this. The lower boundary *q* is in practice also small enough to make the area between *−∞* and *q* be negligible. The general conclusion that 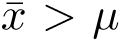 is already stated in [19]. Here, we also observe that 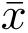 increases as the constraint *x ≥* 2(*r − s*) increases.

Similar to above, let *σ_gd_* be the standard deviation of the distribution of reads spanning a position *p*. Using the relation *xf*(*x*) = *μf*(*x*) − σ^2^ *f*′(*x*), we have

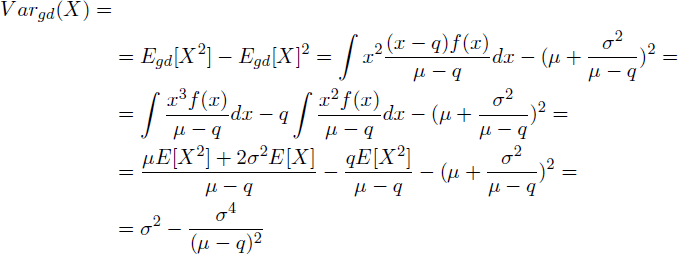

From this derivation. We immediately get Result 2. The approximation in Result 2 is following from the same assumptions as in the derivation of *μ_gd_* above and is a special case of the result in [20]. For hypotheses testing of variants with the assumptions above, *μ_gd_* should be used in *H*_0_ and *σ_gd_* in the significance test.

